# Feasibility in MacArthur’s Consumer-Resource Model

**DOI:** 10.1101/2023.04.14.536895

**Authors:** Andrea Aparicio, Tong Wang, Serguei Saavedra, Yang-Yu Liu

## Abstract

Finding the conditions that ensure the survival of species has occupied ecologists for decades. Theoretically, for mechanistic models such as MacArthur’s consumer-resource model, most of the efforts have concentrated on proving the stability of an equilibrium assuming that it is feasible, but overlooking the conditions that ensure its feasibility. Here we address this gap by finding the range of conditions that lead to a feasible equilibrium of MacArthur’s consumer-resource model and study how changes in the system’s structural and parametric properties affect those ranges. We characterize the relationship between the loss of feasibility and the increase in complexity (measured by the system’s richness and connectance) by a power law that can be extended to random competition matrices. Focusing on the pool of consumers, we find that while the feasibility of the entire system decreases with the size of the pool, the expected fraction of feasible consumers increases —safety in consumer numbers. Focusing on the pool of resources, we find that if resources grow linearly, the larger the pool of resources, the lower the feasibility of the system and the expected fraction of feasible consumers —danger in resource numbers. However, if resources grow logistically, this pattern is reversed with a sublinear increase in feasibility, as it has been previously reported in experimental work. This work provides testable predictions for consumer-resource systems and is a gateway to exploring feasibility in other mechanistic models.

## 1 Introduction

A recurring theme in ecology is identifying the conditions that enable ecosystems to survive. Often, the persistence of a community has been studied using two different approaches. The first approach tries to determine the properties that allow an ecosystem to maintain a long-term constant abundance, or equilibrium, which is related to the system’s *stability*;^1–5^ its robustness characterizes the size of the perturbations that the system can tolerate without drifting away from said equilibrium. One of the best-known results in terms of characterizing the stability of ecosystems is that for a random community whose dynamics close to the equilibrium are described by random pairwise interactions between species, the likelihood of being stable decreases as the complexity increases.^1^ Here, the complexity of the system is measured by the diversity of its species and their interactions.

The second approach tries to determine the ability of the community’s members to coexist, or maintain a positive abundance, described by the concept of *feasibility*.^6–9^ Logically, feasible equilibria are of interest when studying ecosystems and thus, feasibility is considered a necessary condition for persistence. One of the most widely used models in ecology is the Generalized Lotka-Volterra (GLV) model, which captures the community’s dynamics by explicitly modeling the growth rate of its species and the pair-wise inter-species interactions. The stability of the GLV model has been intensively investigated and several Lyapunov functions were proposed that guarantee the global asymptotic stability, provided that the equilibrium point is feasible, for an antisymmetric,^10^ or diagonally stable^11–13^ interaction matrix. Subsequently, the existence of a non-negative and stable equilibrium point for a diagonally stable interaction matrix was proven via a Lyapunov function.^14^ These results formalize the intuition that feasibility is a necessary condition for persistence.

Efforts to characterize the parameter range that leads to feasible equilibria, which we call *feasible parameters*, have been made recently. Notably, Grilli et al. proposed a geometrical interpretation of the range of feasible growth rates, which are contained inside a multidimensional cone defined by the system’s interaction strengths (assumed known); this cone is called the *feasibility domain*.^7, 8^ Further, Song et al. provided a guideline to systematically build the feasibility domain for the GLV model, which includes the case when interspecific interactions are not necessarily static.^15^ This methodology has been used to explore the variable likelihood of different parameterizations of the GLV growth rates leading to feasible, and therefore stable, equilibria for small random communities.^16^ Additionally, an inverse correlation between complexity and feasibility was found when the GLV growth rates were set to the central vector in the feasibility domain.^17^

The GLV model has been an important tool to set the basis for establishing theoretical properties of ecosystems, however, the change of species abundance in nature does not only depend on their pairwise interactions, but also on how individuals enter or exit the community. Beyond the GLV model, other model frameworks that not only include the dynamics of species, but also explicitly model the dynamics of the resources that they consume have been proposed. The most notable of these models is MacArthur’s consumer-resource model^18, 19^ (or MacArthur’s model in short hereafter), in which the interactions between species are represented through the species’ consumption patterns instead of being explicitly modeled. In its original formulation, MacArthur’s model considers that the resources converge quickly to an equilibrium using different timescales. With this assumption, a minimization principle can be applied to demonstrate the global stability of the system given that it is feasible,^18–20^ and the result can be extended to a more general model in the context of niche theory.^21^ Further, the global asymptotic stability of a feasible equilibrium when species and resources’ dynamics evolve in the same timescale have been proved by means of a Lyapunov function which, in fact, can also be used for the GLV model.^22^

Consumer-resource dynamics have been used to model a plethora of communities of many types and sizes,^23^ and several studies generalize the model to include interaction types other than consumption and different resources’ growth rates.^21, 24^ These models are outstandingly helpful when describing the dynamics of microbial communities, where the abundance of the resources that microbes consume is believed to primarily mediate the species’ interactions, so tracking it is crucial to understand the communities’ behavior. Several adaptations of MacArthur’s model have appeared that include different aspects of microbial dynamics. For example, by explicitly including metabolically mediated cross-feeding,^21, 25^ or by assuming that resources interact through chemical mediators including facilitation and inhibition interactions.^26^

In terms of the analysis of coexistence conditions, two special cases of MacArthur’s model have been used to study the feasibility of micro and macroscopic communities. If resources are assumed abiotic and supplied at a constant rate, the feasibility of microbial communities was found to imply its stability.^27^ When the maximum growth rate of resources is equal to their carrying capacity, the seasonal patterns of community composition and interactions between the species can modulate the volume of the feasibility domain of a vertebrate community.^28^ Notably, an experimental setup demonstrated that multispecies microbial communities are possible in cultures where a single resource is provided (a phenomenon attributed to crossfeeding), and that the diversity of the community increases very slowly when the poolk of resources is expanded.^29^

While the stability of MacArthur’s model is very well characterized, we notice a gap in systematically determining the parametric conditions that lead a community to a feasible equilibrium. Here we built, following the geometrical interpretation described for the GLV model, the feasibility domain for MacArthur’s model specifically. We considered two regimes for the resource dynamics: linear and logistic growth. Then, we systematically explored how changes in structural (i.e., properties of network topology such as connectance or nestedness) and parametric (such as niche overlap) properties of the system impact the volume of the feasibility domain of random communities, representing the likelihood of coexistence of all the species. Additionally, we calculate the feasibility volume per species, which represents the likelihood of a single species surviving in its particular community, and can be interpreted as the fraction of feasible consumers; we analyze the impact of the system’s size and connectance on it. We found that the complexity of a system governed by MacArthur’s model is the major driver of the loss of the community’s feasibility. Here, we define complexity as the product between the species richness and the connectance of the underlying network. The degree of its influence depends on the resources’ growth regime and the particular community’s interactions. We characterized the relationship between complexity and feasibility volume through a simple equation employing the physical principle of scaling laws and showed how it extends to competitive GLV systems. Furthermore, we showed that the richness-feasibility trend is reversed for individual species. Because feasibility is a necessary condition for the persistence of the community, a sufficient condition for the stability of MacArthur’s model, and unfeasible states are in general not of interest from a biological point of view, the feasibility domain that we built here can also be understood as a stability domain. That is, the parameters inside it guarantee the community’s coexistence and persistence.

## 2 MacArthur’s consumer-resource model

To model the abundance change of n species and m resources, we consider MacArthur’s consumer-resource model^18, 19^ described by the following set of ordinary differential equations (ODEs):

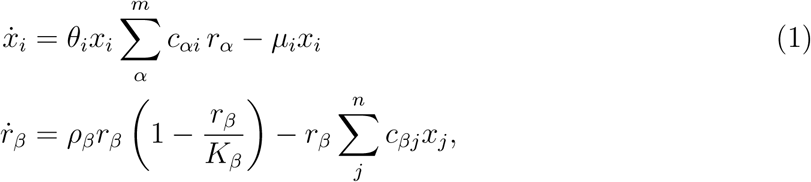

where i = 1, …, n, β = 1, …, m, x*_i_* is the abundance of species i, r*_β_* is the abundance of resource β, θ*_i_ ≥* 0 is the efficiency with which species *i* converts the consumed resources into biomass, and *c_αi_ ≥* 0 is the rate at which species *i* consumes resource *α*. The remaining terms in (1) represent the rates at which species and resources exit and enter the community; µ*_i_* is the mortality rate of species i, and ρ*_β_* and K*_β_* are the maximal growth rate and carrying capacity of resource β, respectively. The vectors of species and resources abundances are given by X *∈* ℝ*^n^* and R *∈* ℝ*^m^*, respectively, while θ *∈* ℝ*^n^*, µ *∈* ℝ*^n^* and C *∈* ℝ*^m×n^* are the vector of species efficiency, the vector of species mortality, and the consumption matrix, respectively. The zero/non-zero pattern of matrix C represents the incidence matrix of a bipartite graph connecting species to the resources they can consume. We will refer to this graph as the *underlying network* of the community (Figure 1.a), where the link direction reflects the flow of energy from the consumed resources to the species.

**Figure 1:**
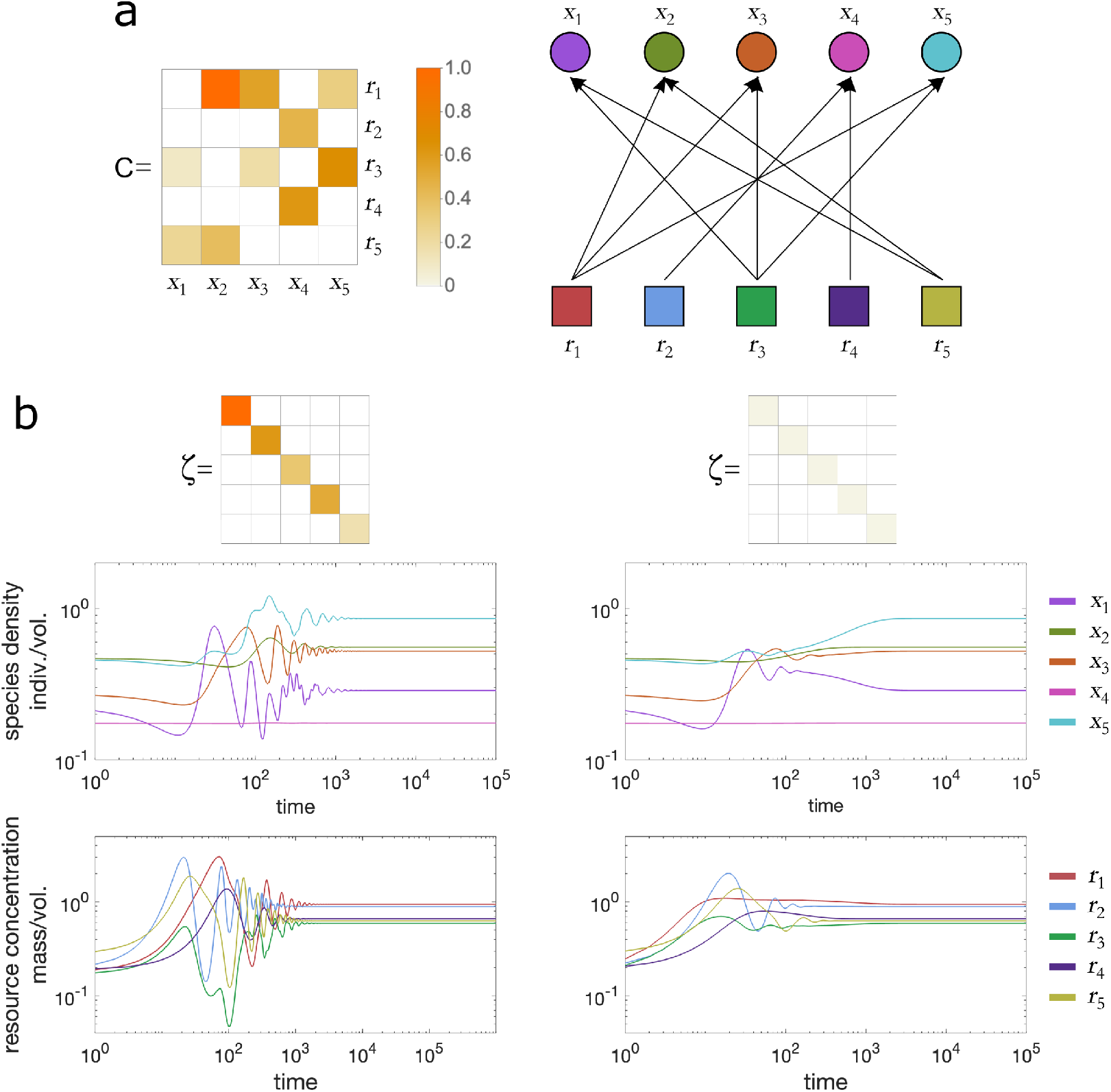
In consumer-resource systems (such as MacArthur’s model) the dynamics of both the species and the resources are explicitly modeled and interactions between species are implicit. **a.** Bipartite underlying network of a consumer-resource system. Here, species X consume resources R and use the consumed biomass to grow. The species’ preference for the different resources is encoded in the matrix C *∈* ℝ^(m×n)^ for n resources and m species. The element *c_βi_* represents the rate at which species x*_i_* consumes resource r*_β_*. **b.** The parameter ζ determines the resource growth, which affects the dynamics of the system. On the left panels, the non-zero elements of ζ are in the same order of magnitude as the consumption rates and the resources grow at a logistic rate. On the right panels, the elements of ζ are close to zero, the resources grow at an approximately linear rate.

## 3 Feasibility domain of MacArthur’s model

The equilibrium solution (X*^∗^*, R*^∗^*) of system (1) is found by solving

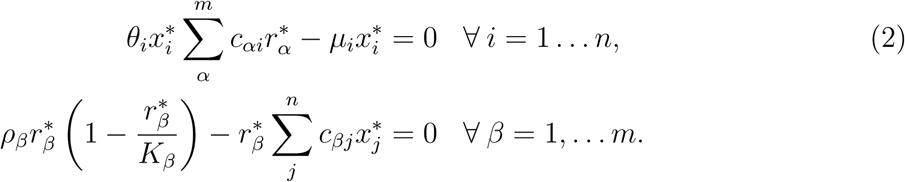

Because negative abundances of species and resources have no biological meaning, and a zero abundance represents an extinction, an equilibrium point with all species surviving is called feasible if a solution to (2) exists such that X*^∗^* > 0 and R*^∗^* > 0, i.e., a nontrivial, strictly positive equilibrium solution. Further, we consider two additional constraints that ensure the biological significance of a feasible equilibrium: i) the mortality rate of any species must have a negative impact on its growth, i.e., *−*µ*_i_* < 0 *∀ i* (or µ*_i_* > 0 *∀* i); ii) the maximal growth rate of any resource must have a positive value, i.e., ρ*_β_* > 0 *∀* β.

MacArthur’s model (1) can be written in the GLV form as^22^

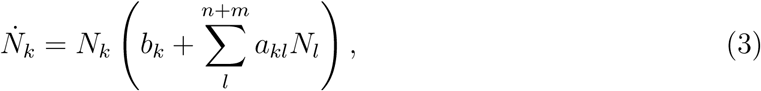

wher

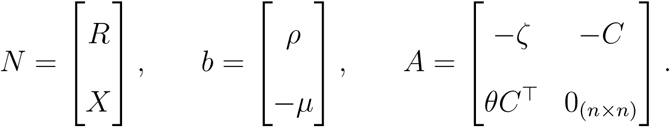

Here, ζ *∈*ℝ^(m×m)^ is a diagonal matrix with elements 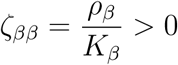.

Clearly, the feasible equilibrium point for (3) is N *^∗^* > 0 which, if A is non-singular, is unique and satisfies

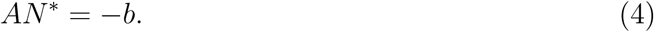

Because ζ is a diagonal matrix, the non-singularity of A depends on C being of full rank, which is satisfied with the assumption that no two species have consumption patterns proportional to each other.

Note that the elements of ζ represent the coefficients for the quadratic part of the resources’ logistic growth, and their numerical values are given by ζ*_ββ_* = ρ*_β_*/K*_β_*. Here, instead of requiring an exact knowledge of the values of ζ, we consider two general cases: the diagonal elements of ζ have a very small value, i.e., ζ*_ββ_ ≈* 0 for all β = 1, …, m; and the diagonal elements of ζ*_ββ_* are within the same order of magnitude as the elements in the corresponding row of C. These two cases represent two distinct dynamic regimes. In a), the growth rate of resource β is,

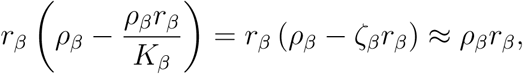

which is indicative of its carrying capacity being much larger than its maximal growth rate. This can occur in communities where resources are often depleted to very low concentrations as long as they are consumed by some species, but if the consumption ceases, the concentration would increase largely. For example, this might be the case of microbial communities with self-renewing nutrients. The left panels in Figure 1.b show the dynamics of this case. In case b), the resources grow at a logistic rate, i.e., they grow fast at small abundances, and the growth stops when the abundance reaches the carrying capacity. This case is observed in large-scale natural ecosystems such as plants-animals networks. The right panels in Figure 1.b show the dynamics of case b). In the following, we will focus on case a) while drawing parallelisms with case b).

For any given non-negative species abundances and resource concentrations, i.e., N *^∗^* = [X*^∗^* > 0 R*^∗^* > 0]*^⊤^*, it is evident from (4) that a vector b = [ρ, *−*µ]*^⊤^* that satisfies the constraints i) and ii), with ζ*_ββ_* > 0 and *c_αi_ ≥* 0 for all (β, *α*) = 1, …, m, and *i* = 1, … n, can always be found. On the other hand, characterizing the values of b and A that will lead to a feasible equilibrium, i.e.,

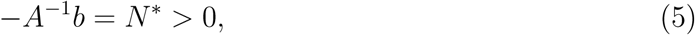

is a more interesting and harder task. For a given A only certain values of b, i.e., some sets of maximal growth rates ρ and mortality rates µ, will satisfy (5). We call these values of b *feasible parameters*; each unique set of feasible parameters will take the system to a different equilibrium (X*^∗^*, R*^∗^*). Equation (5) implies that a necessary condition for the feasible coexistence can be determined by the interplay between the consumption pattern of the species and the rates at which the resources and species enter and exit the community, respectively. The portion of the S-dimensional space (S = n + m) where all the possible feasible parameters for a given community live is known as the *feasibility domain*.^7^ It has been proven that any feasible equilibrium point of the consumer-resource model in the form (3) is globally and asymptotically stable.^22^ Therefore, the feasibility domain captures all the parameters that can generate communities with all positive abundances by numerically solving the ODEs of MacArthur’s model. This framework is directly related to the feasibility formulation for GLV systems (See SI 1) for details on the feasibility domain of the GLV model). Hence, in the following, we will draw parallelisms between the feasibility of MacArthur’s model and competitive GLV models (where matrix A is a square matrix with negative elements and its zero/non-zero pattern is symmetric).

The feasibility domain has a very intuitive geometrical interpretation. To get a sense of its construction, let’s consider the toy community in Figure 2.a with one species consuming one resource (S = 2), whose dynamics are governed by system (1). Assume that the consumption rate c, and the parameters θ and ζ are constant and known, such that writing the dynamics in the GLV form yields a matrix *A ∈* ℝ^2×2^. From (4), for S = 2, we can write 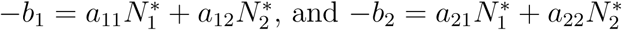, which indicates that the columns of A contain information to determine the possible feasible values of b. By normalizing the columns of A we can obtain the unit vectors (or *generating vectors*) g_1_ and g_2_ that have a start point at the origin of a 2-dimensional space, and their endpoints intersect the unit circle centered at the origin (top right panel of Figure 2.a). The axis of this 2-dimensional space are labeled (ρ, µ) and the coordinates of every point contained in the convex cone generated by vectors g_1_ and g_2_ represents a pair of feasible parameters for our toy community. Naturally, this procedure can be generalized for S dimensions, which allows us to find the convex cone containing the feasibility domain of any system in the form (3) (Figure 2.b shows the feasibility domain construction for a community of size S = 3 with n = 1 species and m = 2 resources). This procedure has been used, and described in detail, several times in the literature.^9, 15, 16, 30^

**Figure 2:**
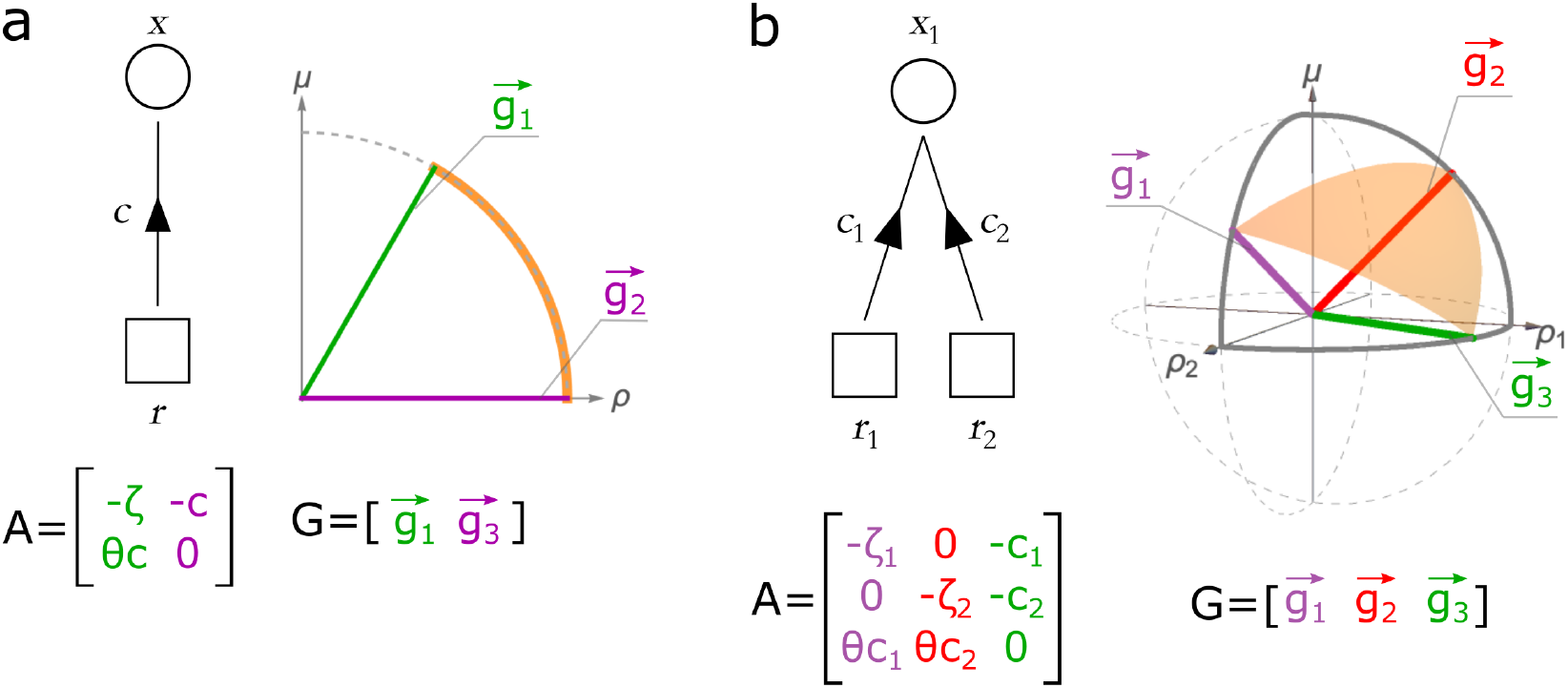
MacArthur’s model with n species and m resources can be written in the GLV form *N* = *N*_diag_ (*b+AN*), where *N* =, and A *∈* ℝ(*^S×S^*) is interpreted as an adjacency matrix with ζ*_β_* = ρ*_β_*/K*_β_* and where the system’s dimension is S = n + m. The normalized columns of A generate S unit vectors ⃗g arranged in matrix G. These vectors delimit a polyhedral cone in a S-dimensional space that contains the feasibility domain. **a.** For the simplest community of one species consuming one resource (S = 2), the normalized columns of A *∈* ℝ^(2×2)^ (green and purple in the panel) generate two unit vectors 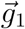 and 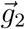 (colored accordingly) that span a cone in the two-dimensional space (ρ, µ). This cone contains the system’s feasibility domain, whose size is quantified as the fraction of the unit circle that is intersected by the feasibility domain (orange arch). **b.** For a community of size S = 3 that has one species consuming two resources, the normalized three columns of A (purple, red and green) generate three vectors that span a volume in the three-dimensional space (ρ_1_, ρ_2_, µ). The fraction of the unit sphere that the cone’s solid angle (orange surface) intersects quantifies the size of the system’s feasibility domain. Because the growth and mortality rates must be positive in order to maintain their biological significance, the maximal size of the community’s feasibility domain is Ω = 0.25, i.e., one quarter of the unit sphere (delimited by the solid gray arches).

Because a system’s feasibility domain contains all its possible feasible parameters, we can interpret its size as a proxy of the probability of randomly sampling feasible vectors of maximal growth and mortality rates. In other words, the larger a community’s feasibility domain, the higher the probability that all its members will survive. It follows that the feasibility domain’s size is equivalent to its corresponding convex cone’s volumetric modulus, loosely defined as the intersection of the cone’s S-dimensional volume, and a closed unit ball in R*^S^* (see SI 2 for details). The orange arch in Figure 2.a and the orange surface in Figure 2.b show the volumetric modules for our 2 and 3-dimensional examples, respectively. Following well established methodologies,^31^ the volumetric modulus of a convex cone can be calculated yielding, for this application, a numerical value for the probability of survival of a community (see SI 2.1 for details). We will refer to the volumetric modulus as the *volume of the feasibility domain*, denoted as Ω hereafter. Note that Ω represents the probability of survival of the whole community, that is, of all the species having a positive abundance at equilibrium. Looking at the species level, we can calculate the feasibility per population ω = Ω^1^*^/n^*,^32^ and interpret it as the probability of feasibility of a single species.^16, 33, 34^ In other words, ω represents the probability of survival of each individual species in its particular community, or the expected fraction of feasible consumers. We will refer to this quantity as the *feasibility per species*.

Due to the constraints µ*_i_* > 0 and ρ*_β_* > 0, the maximal volume of system’s (1) feasibility domain is of one quarter of the unit ball (Ω = 0.25), i.e., only the positive side of the µ and ρ axes can be occupied by it. The maximal feasibility domain for the size S = 3 example is delimited by the solid gray arches in the top-left of Figure 2.b. A small Ω thus implies that a small parametric perturbation could potentially lead to the loss of coexistence. On the other hand, the maximal Ω = 0.25 implies that the probability of randomly sampling feasible vectors of maximal growth and mortality rates equals one so that no parametric perturbation can lead to an unfeasible equilibrium. In this sense, the size of the feasibility domain can be understood as a type of robustness.

## 4 Results

We aim to understand the effect of parameter variations on the feasibility of a consumer-resource community, and how the dynamics of MacArthur’s model influence its feasibility compared to that of competitive GLV. To achieve that, we generated random communities of different sizes S and we systematically changed their underlying network connectance (κ) measuring the feasibility volume Ω at every step. Communities governed by MacArthur’s model, have a bipartite underlying network of size S = n + m (we initially assumed n = m and then analyzed the case of n *̸*= m), while communities governed by competitive GLV have an undirected network of size S. We parameterized MacArthur’s model’s consumption matrix C, efficiency vector θ, and diagonal elements of the resources growth rate ζ by drawing random numbers from normal distributions *N* (0, σ), *N* (0, σ*_θ_*), and *N* (0, σ*_ζ_*) respectively, and then taking the absolute values. We parameterized the competitive GLV’s interactions matrix A by drawing random numbers from *−*abs(*N* (0, σ)) (details in SI 3). Additionally, we investigated the impact of nestedness (a structural property) and niche overlap (a parametric property) on the feasibility volume of consumer-resource communities.

### 4.1 Larger and more connected communities have a smaller chance of coexistence

Because we assume a consumption matrix C of full rank with n = m, the minimal connectance of the community’s underlying network is κ = n/(nm), i.e., every species is connected to only one resource and every resource is connected to only one species (all species consume *exclusive* resources). In other words, minimal connectance implies that species do not compete for resources at all. In this case, as long as the mortality and maximal growth rates satisfy the conditions µ > 0 and ρ > 0 (conditions i) and ii)), Eq (5) readily indicates that the equilibrium is guaranteed to be feasible, i.e., the feasibility volume is maximal Ω = 0.25 (SI 2). We generated communities with minimal connectance, random consumption rates and very small ζ, and we gradually added random interactions, thus increasing their connectance κ (details in SI 3), calculating their feasibility volume Ω and the feasibility per species ω = Ω^1^*^/n^* at every step. We found a monotonic decrease in Ω as connectance κ increases (Figure 3.a shows the feasibility volume of 1000 communities of size S = 6 (green crosses), S = 10 (pink triangles), and S = 20 (blue diamonds) see Figure S4 for other community sizes). This implies that as species consumption patterns become more similar, small parametric perturbations are more likely to lead to a loss of feasibility. Note that an increase in connectance can be interpreted as the species increasingly sharing consumption preferences or, in other words, an increase in competition for resources. We also found that the feasibility volume Ω of larger communities tends to decrease faster as κ increases, than that of smaller ones, indicating that larger communities are less likely to survive as a whole. Interestingly, the feasibility per species tends to grow with the community size, but still decreases with increasing connectance (Figure 3.b). This means that individual species find safety in numbers and are more likely to survive in larger communities. Furthermore, these two results extend with no qualitative changes to random competition systems governed by GLV dynamics (Figure 3c-d), and are robust to the use of different distributions to obtain the random parameters (SI 4)). Remarkably, for MacArthur’s model when ζ*_ββ_* takes values in the same order of magnitude as *c_αi_*, while the general trends are maintained in terms of whole community survival, the connectance-volume curve flattens substantially at low connectance (Figure S5.a). This is explained by the influence that the non-negligible quadratic part of the resources’ growth rate has on the feasibility domain’s generating vectors. More precisely, at low connectance, the first m normalized columns of A have similar weights in the first n elements (given by ζ) and the last n elements (given by θC*^⊤^*), pulling the generating vectors away from the ρ axis in the parameter space. As connectance of C increases, the consumption interactions represent the majority of the weights in the generating vectors (Figure S6). In terms of feasibility per species, there are no qualitative changes induced by the magnitude of ζ (Figure S5.b).

**Figure 3:**
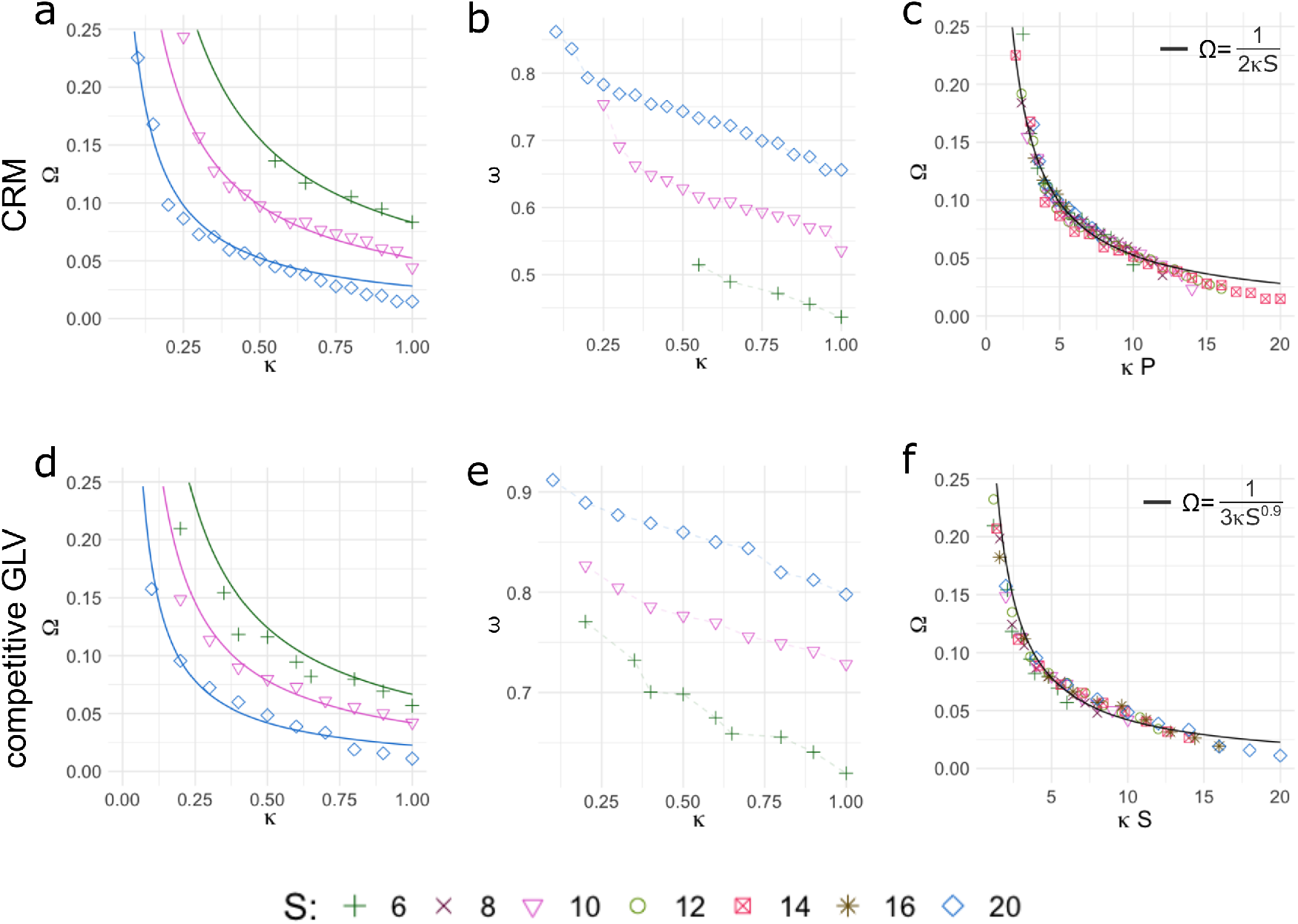
Increasing complexity leads to a monotonic loss of feasibility robustness for consumer-resource (panels **a**-**c**) communities with linear growth, and competitive GLV communities (panels **d**-**f**) We generated communities of different sizes and randomly added interactions between its members (consumption interactions for MacArthur’s model and competitive interaction for the GLV), which varies the underlying network connectance. We measured the size of the resulting feasibility domain at every connectance change. The interaction strengths and efficiency factors were sampled using the absolute value of normal distribution with zero mean and constant standard deviation. **a**. Volume of the feasibility domain Ω of communities of various sizes S (here S = n + m and n = m) with ζ 0, with respect to the community’s connectance κ. As species consumption patterns become structurally more similar (larger κ) and communities become richer (larger S), the feasibility volume monotonically decreases, and so does the probability of randomly sampling parameters that lead to the whole community’s survival. **b**. Feasibility per species (ω = Ω^(1^*^/n^*^)^), with respect to the community’s connectance κ (horizontal axis). Individual species’ feasibility decreases with an increased connectance but increases with community size. Individual species have a larger probability of survival when they are a member of richer communities. **c**. The product κS (directly proportional to the communities’ complexity) is a scaling variable for the feasibility volume of communities of various sizes and connectances. This results in a data collapse that unveils the complexity-feasibility volume relationship governed by the power law 1/Ω = a(κS)*^q^*. **a,c**. The solid lines trace the complexity-feasibility function with a = 0.5 and p = 1, tracking the data obtained from our simulations very closely. **d**-**f**. The loss of feasibility with increasing complexity trend, and increase of feasibility per species with richness, is extended to systems with general competition matrices.

### 4.2 The likelihood of feasibility is inversely proportional to the system’s complexity

We now investigated the interaction of connectance and community size and its impact on the feasibility volume. We found that using the community size as a scaling factor for the Ω vs. κ relationship, in communities governed both by MacArthur’s model with small ζ and competitive GLV dynamics, leads to the data collapse shown by the markers in Figure 3.c and 3.e, respectively (markers represent the mean Ω of 1000 communities of sizes that range from S = 6 to S = 20, and different connectances). This suggests the existence of a universal relationship between κS and Ω, for any community size, that is robust to different parameterizations (Figure S1). Therefore, we can define κS as a measure of the system’s complexity and conclude that the community is less feasible when κS is larger. Indeed, by performing a linear regression on the log *−* log transformation of the data (Figure S7) we can estimate the parameters of a power law of the form y = ax*^b^* (see SI 5 for details) where the independent variable is x = κS and the dependent variable is the inverse of the feasibility volume y = 1/Ω. For instance, the best fit for our examples of MacArthur’s model and of competitive GLV are a = 2, b = 1 and a = 3, b = 0.9, respectively. This unveils the simple relationship between the feasibility volume and the complexity of the community given by

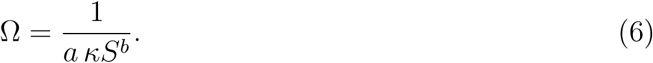

The solid lines in Figure 3.a-c show the curve (6) for the a range of the communities’ connectances and richness (other community sizes in Fig S8). All the curves follow very closely the data obtained from our simulations so we conclude that the community’s complexity is a strong driver of the feasibility loss. It is surprising to note that our defined complexity is highly analogous to the definition raised by Robert May 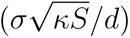^1^ to explain the local stability of a feasible solution for systems with random interaction matrices. The fact that the product between κ and S similarly contributes to the system’s complexity in these independent studies with different focuses shows its importance when analyzing the persistence of a large ecosystem. Because connectance is less relevant for the feasibility probability when ζ*_ββ_* is in the same order of magnitude as the non-zero *c_αi_*, especially at low connectances, it is no surprise that the complexity-feasibility relationship does not readily extend to this case Figure S9.a. However, because higher connectance leads to a higher influence on the feasibility domain, and richness maintains its importance at any value of ζ, a power law that describes the complexity-feasibility relationship can be found when κ *≥* 0.4 (Figure S9.b). We conclude that complexity is the main driver of feasibility when the interactions between species (for the competitive GLV) or between species and resources (for MacArthur’s model) dominate in the community’s dynamics.

### 4.3 The resources growth dynamics determine the complexity-feasibility relationship when **n < m**

To analyze the influence of connectance and richness on the feasibility of communities governed by MacArthur’s model with more resources than species, we generated random communities with a fixed number of species (n = 5) and a varying number of resources (m = [5, 9]), and gradually varied the number of consumption interactions between them (modifying κ). Again, we considered the two cases of resources growth rate, determined by the magnitude of the diagonal elements in ζ: a) ζ*_ββ_ ≈* 0, i.e., the resources grow at an approximately linear rate, and b) ζ*_ββ_* is in the same order of magnitude as the consumption rates in C, i.e., the resources grow at a logistic rate with a maximum imposed by their carrying capacity.

In case a), we observed a qualitatively equivalent connectance-feasibility trend to the case of n = m, i.e., the feasibility volume and feasibility per species decrease as communities become more connected (Figure 4.a-b). Additionally, the feasibility volume decreases with community richness as well, maintaining the complexity-feasibility relationship. However, a larger pool of resources leads to smaller feasibility per species, reversing the “safety in numbers” trend that we had observed when n = m and making every individual species less likely to survive when the number of resources expands (Figure 4.b). In case b), an increasing connectance remains the driver of the loss of feasibility in the whole community, but the complexity-feasibility relationship found in n = m doesn’t hold because the expansion of the resources pool leads to an increase in the feasibility probability (Figure 4.c). This behavior is what one could intuitively expect in large-scale communities where self-renewable resources grow at a logistic rate capped by a carrying capacity, when species have more resources to choose from. Indeed, the feasibility per species in Figure 4.d shows that individual species also have a better chance of survival in communities where resources are more diverse than in communities with fewer resources.

**Figure 4:**
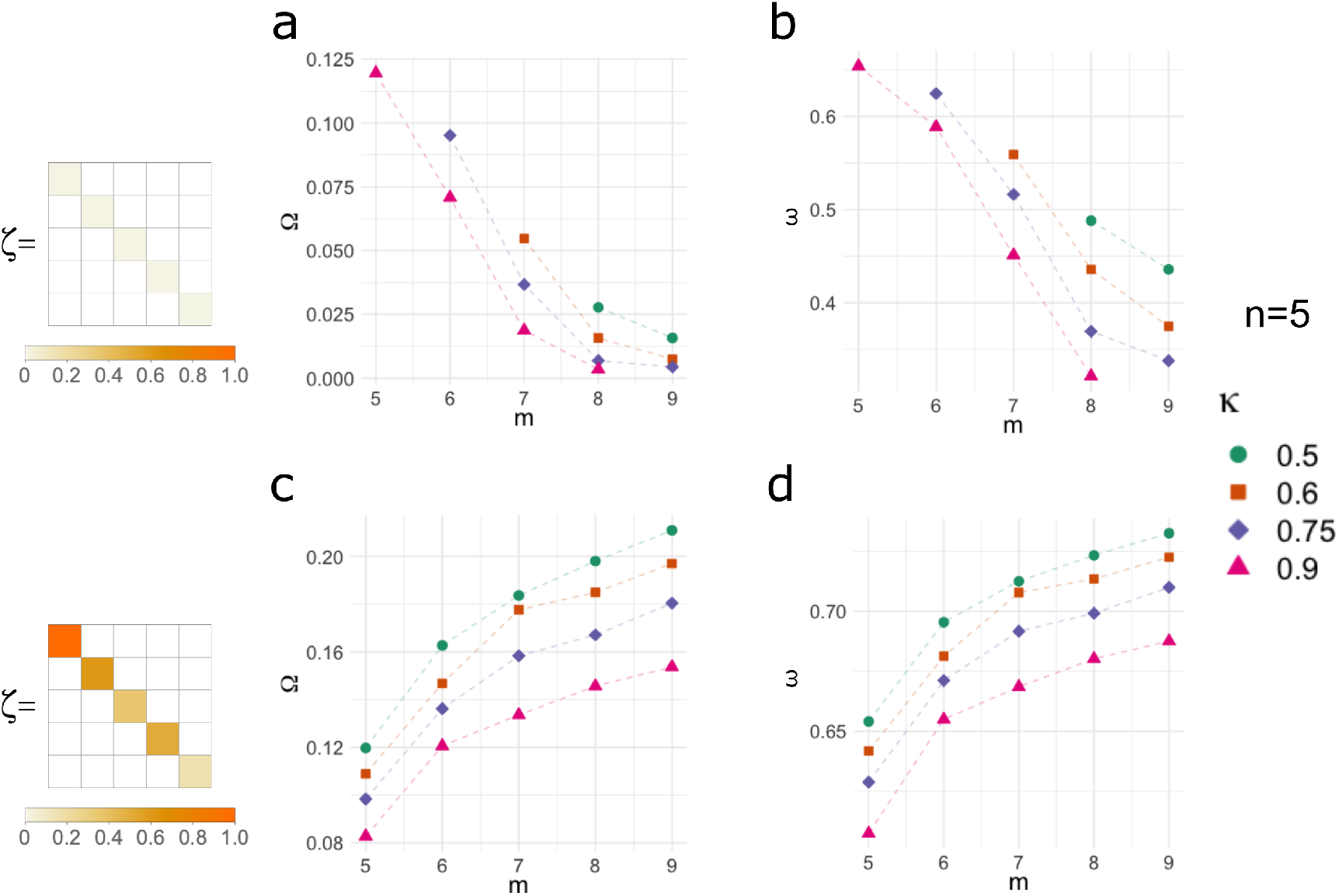
Feasibility volume and feasibility volume per species for communities governed by MacArthur’s model with a fixed number of species (n = 5) and a varying number of resources (m = [5, 9]) Here, the growth rate of resources determines the complexity-feasibility relationship. **a** and **b**. When resources grow linearly (ζ ≈ 0), communities that are more interconnected (i.e., larger κ) and have a larger number of resources, tend to have a smaller probability of feasibility. **b.** Contrary to the case of n = m and rather counterintuitively, the feasibility per species decreases as the number of available resources grows. **c** and **d**. When resources grow at a logistic rate (ζ in the same order of magnitude as the consumption rates), communities with a larger resource pool tend to have a larger probability of being feasible, while increasing connectance remains a driver of the loss of feasibility. **c**. The feasibility volume tends to grow with the resources pool, reversing the trend observed when n = m and increasing the probability of the whole community to survive. **d**. The feasibility per species also tends to increase as the pool of species grows, which is intuitively expected for larger scale communities whose resources grow following a logistic rate.

### 4.4 Increased nestedness leads to a loss of feasibility

Next, we analyzed the impact of nestedness on the feasibility volume. Nestedness is an ecological pattern that occurs when specialist species tend to interact with resources that are consumed by more generalist species^35^ (the top panel in Figure 5.a shows the nested structure of the consumption matrix C and the underlying bipartite network). This type of structure has been observed in many empirical ecosystems, and reported to decrease competition, facilitate species coexistence,^36^ and promote the community’s persistence.^9^

**Figure 5:**
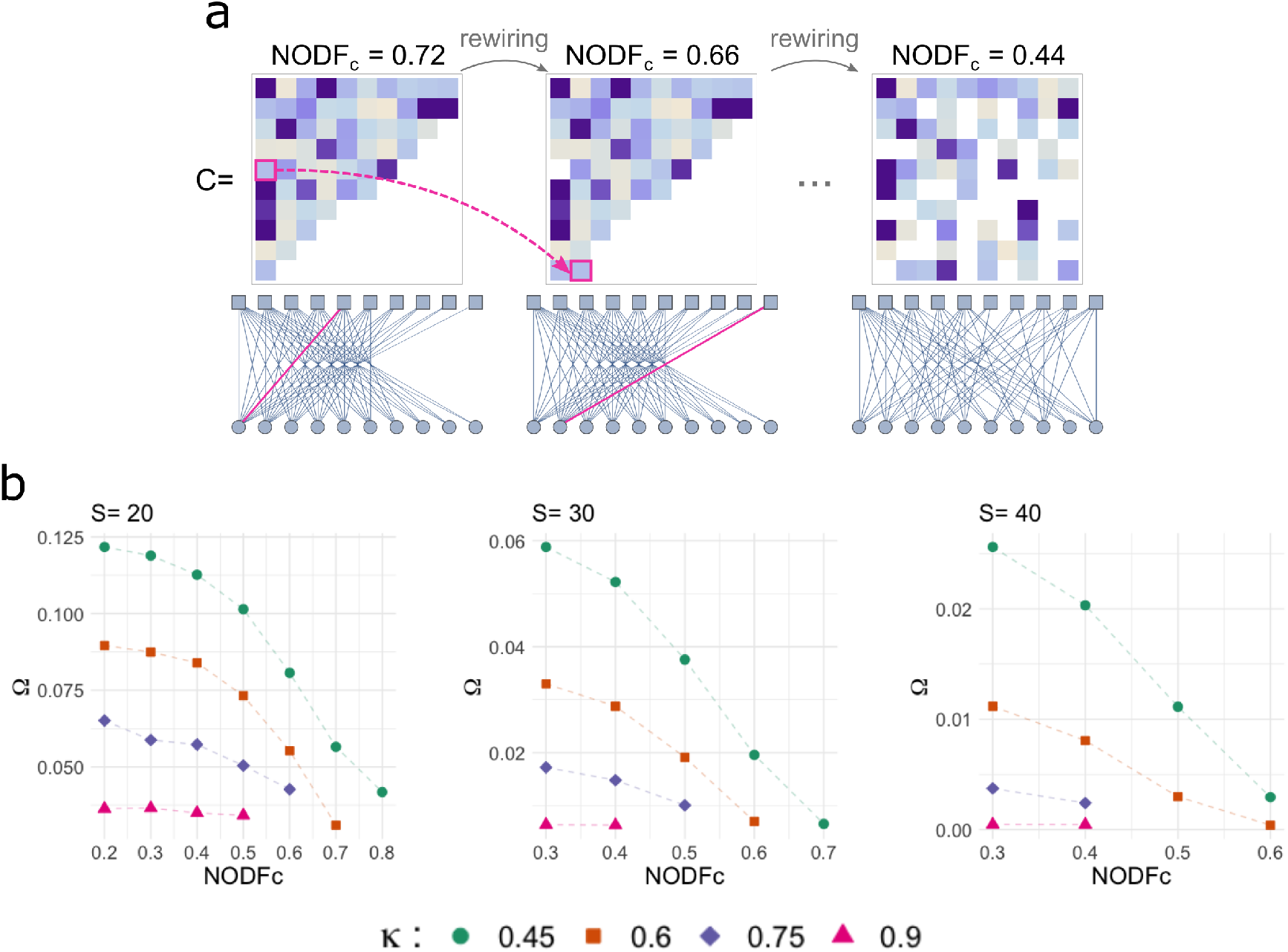
More nested communities tend to have a smaller feasibility domain. In nested communities, specialist species tend to interact with resources that are consumed by more generalist species; nestedness is a structural property of the underlying network. **a.** We quantify the nestedness using the nestedness metric based on the overlap and decreasing fill represented by NODF = (0, 1] where 1 is perfect nestedness, and normalize it controlling for connectance and richness as 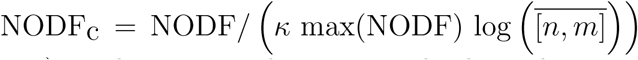. For a community of S = 20 (n = m = 10), and κ = 0.6, the top panels show the consumption matrix C and the bottom panels show the corresponding bipartite underlying graph where species are circles and resources are squares. The left panels show the community structure with the maximum NODF_c_, the center panel shows one step of the rewiring process by which an interaction between a species and a resource is relocated to another pair that didn’t interact previously (pink squares on C and pink links in the underlying graph). We iteratively performed this rewiring procedure to decrease the nestedness (measured by NODF) while preserving the connectance and the distribution of interaction strengths. The right panel shows the result of iterating 10 rewiring steps, only accepting the rewirings that decrease the NODF_c_ and rejecting those that don’t. **c.** Increased nestedness, in addition to increased connectance, is a driver for the loss of feasibility. The panels show the feasibility volume of communities of size S = (20, 30, 40) with respect to their NODF_c_ score (horizontal axis), for several connectance values (colored markers). We found that for a given connectance, increasing the NODF_c_ tends to result in a smaller feasibility domain, especially for less connected communities.

We quantified nestedness using the overlap and decreasing fill (NODF) measure^35^ which can take values between zero and one; NODF=1 is the nestedness score for a perfectly nested community (left panel in Figure 5.a and SI 6). To adequately compare nestedness across different networks and correct for correlations with connectance and richness,^37, 38^ the NODF measure is normalized as 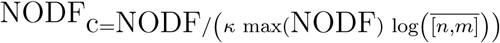,^38^ where max(NODF) is the maximum possible NODF for a network of a specific connectance, and 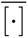 denotes the geometric mean (see SI 6.1 for details).

To test the influence of nestedness on the feasibility volume, we generated communities with a structure that has the highest NODF possible for a fixed connectance, and a matrix C of full rank. Again, we parameterized matrix C by drawing random numbers from *N* (0, σ) and taking the absolute values. Then, we ran an edge rewiring algorithm to decrease the NODF while preserving the system’s connectance κ and distribution of interaction strengths (Figure 5.a, see SI 3 and SI 6 for details), and measured the feasibility volume Ω and calculated NODF_c_ with each step. Figure 5.b shows the results of tracking the feasibility volume Ω with respect to the NODF_c_ score for different connectances κ, in communities of size S = (20, 30, 40). We found that Ω generally follows the trend of communities with random structure, i.e., larger κ leads to a smaller Ω, but, interestingly, as the communities become more nested, Ω also tends to decrease monotonically. The decrease is steeper for communities with smaller κ. Therefore, increasing nestedness, along with increasing connectance, are both drivers of a monotonic decrease in feasibility volume.

This might seem to contradict previous reports of increased coexistence in nested communities but, in fact, is directly related to the conclusion for random communities. Recall that any deviation from the case of all specialists that consume exclusive resources decreases the range of feasible parameters or, in other words, any increase in competition leads to a loss of feasibility. Because nestedness leads to a larger overlap of species’ consumption patterns compared to random interactions, due to the competition for some resources becoming extremely strong, it makes sense that more nested communities tend to have a smaller feasibility domain.

### 4.5 Niche overlap leads to loss of feasibility

Additionally, we analyzed how changes in the niche overlap impact the feasibility volume of the community. A species’ niche is generally understood as the range of environmental factors that permit a species to persist,^39, 40^ where those factors include the resources that they consume. Thus, the niche overlap measures the similarity in resource consumption of two species,^40^ although there is no definite consensus on how to quantify it. To vary the similarity of resource consumption between species in a random community with n = m, we first picked a set of n unique pairs of interacting species and resources such that no species or resources are repeated in the pairs. We call these the *preferred interactions* (Figure S3.a). Then, we re-sampled their consumption rates by drawing numbers from a normal distribution with the mean of η_d_ *≥* 0 and standard deviation σ, and then taking the absolute value. We quantified the parametric similarity between the preferred interactions and the rest as

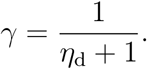

Here, γ is a measure of the niche overlap between species: γ = 1 represents an evenly distributed competition for resources and, as γ decreases, the competition for the species’ favorite resources gets weaker (see SI 7 for more details). We find that for a given connectance, communities with a smaller γ tend to have a larger feasibility domain across the community sizes (Figure S3b). We can conclude that the parametric dissimilarity reduces the niche overlap between species and thus, consistently with previous results,^41^ contributes to a higher likelihood of coexistence.

## 5 Discussion

Consumer-resource models have repeatedly appeared in the ecology literature for the past 50 years. Several variants of the classical MacArthur’s model have been proposed for different specific applications, but always under the same mechanistic framework where inter-species interactions are implicitly given by the depletion of resources in their environment. The global stability of MacArthur’s model has been thoroughly studied and fully characterized in the past, assuming the feasibility of an equilibrium point.

Here we revisit this classical model in its original form (resources grow at a logistic rate and species deplete them, using the consumed biomass to grow), and with a modification on the resources growth rate (they grow at a linear rate without an upper bound). Because for a fixed set of consumption rates only certain values for the mortality and maximal growth rates lead the system to a feasible equilibrium (where all its members survive), we explore how structural changes in the community determine the range of these values. We follow a geometrical interpretation of the feasibility domain^15^ and build a polyhedral convex cone in a multidimensional space that contains all the possible feasible parameters. This technique has been used before to search for the range of feasible growth rates for the GLV model, that model pairwise interactions between species only. Certainly, MacArthur’s model can be written in the form of the GLV and has been mentioned in passing when studying the feasibility properties of the latter. Here, we describe the parallelisms with the general competitive GLV, but also discover important differences that derive from the particular consumer-resource dynamics.

Previous studies have formalized the feasibility of MacArthur’s model’s equilibrium as a sufficient condition for its stability (or persistence)^19, 21, 22^ but, because stable and unfeasible states are not of interest in Ecology, feasibility is also considered a necessary condition. The feasibility of a community represents the ability of its members to coexist, so the size of an ecological system’s feasibility domain is a measure of its robustness, i.e., it characterizes the size of parametric perturbations that the community can endure without losing members to extinction. Here we investigate, via numerical simulations, how changes in the communities’ structural properties can affect the volume of the feasibility domain. Our main result sheds light on the relationship between the system’s complexity and its feasibility domain volume. In particular, we find that the feasibility domain, in general, tends to monotonically shrink when communities are larger and/or more connected. This is congruent with MacArthur’s conclusion on how the stability of ecosystems usually decreases when species are added,^19^ and with results that found a negative correlation between species richness and the likelihood of feasibility in the GLV model.^17^ Moreover, by expressing the community’s complexity as the product between the connectance of its underlying network and its size, a simple expression that closely describes the complexity vs. volume inverse relationship for both MacArthur’s model with linear growth and competitive GLV is unveiled. The complexity-stability relationship in ecological systems has been studied several times before,^5, 17^ usually leading to the conclusion that complexity leads to a loss of stability, but a mathematical expression that models the complexity-feasibility relationship had never been introduced before. Furthermore, our result is consistent with the complexity-stability conclusion because feasibility is a sufficient condition for stability and a necessary one for persistence and addresses the need for investigating the connectance of the community as an independent parameter, explicitly reported in past literature.^17^

While part of our results highlights the similarities between MacArthur’s model and the competitive GLV feasibility properties, we also found that the very particular consumer-resource dynamics induce deviations from the general competition case that justify being studied separately. For example, for systems governed by MacArthur’s model, the product of the connectance and the system’s size is a scaling factor when resources grow linearly, but this property holds only for larger connectances (κ > 0.4 in our example) when resources grow logistically. This reveals that complexity is the main driver of feasibility when the consumer-resource interactions dominate on the dynamics; in the case of low connectances and logistic growth, the resources growth rate has a strong influence over the whole community’s dynamics. When the available resources outnumber the species, the resources grow regime again determines the complexity-feasibility relationship. If resources grow linearly, the increasing complexity-decreasing feasibility trend of the whole community is qualitatively maintained, but the expected fraction of surviving species decreases when the resource pool grows. On the other hand, when resources grow logistically, the whole community and individual species have larger chances of feasibility when pool of resources is larger. More specifically, the fraction of feasible consumers grows sublinearly as the resources become more diverse. This phenomenon had already been observed in an experimental setting and our result provides confirmation of it being the expected outcome when resources are added to a community. Furthermore, increasing connectance remains a driver for the loss of feasibility in all cases.

Additionally, we explored the impact of the community’s nestedness and niche overlap on its feasibility volume Nestedness is a structural property where species’ competition for resources is arranged in a nested manner, while the niche overlap is a parametric property that quantifies the amount of competition that species endure for particular resources. We find that the degree of nestedness and niche overlap in a community have an impact on the size of its feasibility domain but don’t modify the decreasing trend that higher complexity induces. We attribute the inverse relationship between nestedness and niche overlap and the size of the feasibility domain to the increasing competitions that some species face for the resources that they consume. In nested communities, the competition that specialists find overpowers the generalists’ ability to preserve a niche without much competition, especially in communities with a low connectance.

In summary, here we have explored the robustness of a consumer-resource model from the perspective of its equilibrium feasibility. Our results bridge the gap between the assumption of MacArthur’s model equilibrium’s feasibility and the proof of its global stability and show the importance of considering the specific regime under which a community’s resources self-renew when determining its likelihood of coexistence and persistence. Because our results are mostly based on structural properties of MacArthur’s model’s consumption matrix, we hypothesize that similar complexity-feasibility trends will be found when interactions between consumers and resources dominate the dynamics of the community. However, based on the important differences induced by resources growth regime that we observed, we conclude that more research is needed to carefully study the impact that different kinds of interactions might impact the feasibility volume.

## Author contributions

A.A and Y.Y.L conceived and designed the project; A.A did the simulations; all authors analyzed and interpreted the data; A.A. wrote the manuscript with the help of T.W.; all authors revised the manuscript.

