## Supplementary Information for "Feasibility in MacArthur’s Consumer-Resource Model"

### Supplementary Information
### Feasibility in MacArthur's Consumer-Resource Model

Andrea Aparicio[1], Tong Wang[1], Serguei Saavedra[2], and Yang-Yu Liu[1,3]

[1] Channing Division of Network Medicine, Department of Medicine, Brigham and Women's Hospital and Harvard Medical School, Boston, MA 02115, USA

[2] Department of Civil and Environmental Engineering, Massachusetts Institute of Technology, Cambridge, MA 02139, USA

[3] Center for Artificial Intelligence and Modeling, The Carl R. Woese Institute for Genomic Biology, University of Illinois at Urbana-Champaign, Champaign, IL 61801, USA

**SI 4 Robustness with respect to parameter distribution** **8**

**SI 5 Scaling and power laws** **9**

**SI 6 Nestedness** **11**
SI 6.1 NODF normalization and control for modifiers . . . . . . . . . . . . . . . . 13

**SI 7 Niche overlap** **14**

**SI 8 Additional Figures** **17**

**SI 9 References** **20**

### SI 1 Feasibility domain for the GLV model

In any ecosystem, the species biomass or abundance cannot take negative values, and neither can the variables $N$ representing them in mathematical models, without losing biological meaning. The equilibrium of such models $N^*$ must, therefore, be positive. This necessary property of an ecosystem model is called a *feasible equilibrium* [1]. Formally, a feasible equilibrium of a system with $P$ species is defined as any equilibrium that belongs to the interior of the positive orthant of the phase space [2], i.e.,

$$N^* \in \text{ int } \mathbf{R}_+^P = \{x \in \mathbf{R}^P : x_k > 0, \forall\, k = 1, \ldots, P\}.$$

The generalized Lotka-Volterra equations are

$$\dot{N}_k = N_k \left( b_k + \sum_l^P a_{kl} N_l \right),$$

where $b_k$ is the intrinsic rate of growth (increase or decrease) in the abundance of species $k$, and $a_{kl}$ $(l = 1 \ldots P)$ represents the interaction strength of species $l$ on species $k$. The interspecies interactions are gathered in a matrix $A = (a_{kl})$, and the growth rates in a vector $b = [b_1, \ldots, b_P]^\top$. If $A$ is non-singular, the equilibrium $N^*$ is unique and satisfies

$$AN^* = -b. \tag{1}$$

When the vector $b$ is such that $N^*$ lies in the positive orthant of the $P$ dimensional space, i.e., $\mathbf{R}_+^P$, the equilibrium is feasible. The subset of the $P$-dimensional parameter space spanned by all the vectors $b$ that yield a feasible equilibrium is called the *feasibility* *domain* $\Omega$, i.e.

$$\Omega \in \mathbf{R}^P = \{b \in \mathbf{R}^P : -A^{-1}b > 0\}. \tag{2}$$

From (2) it follows that

$$\Omega = \{N_1^* \vec{a}_1 + \cdots + N_P^* \vec{a}_P, \text{ where } N_1^* > 0, \ldots, N_P^* > 0\},$$

where vector $\vec{a}_k$ is the $k$th column of matrix $A$. Geometrically, the feasibility domain is described as a polyhedral convex cone [3] in the $P$-dimensional parameter space whose $k$th border, also called a generating vector [4] or extreme ray, is given by the vector $\vec{a_k}$ [2, 5, 4]. Due to the linearity of (1), if $b \in \Omega$, then $cb \in \Omega$ for any real constant $c > 0$. Thus, without loss of generality, we can focus our analysis to the generating vectors of length 1, given by the normalized columns of $A$, i.e., $\dfrac{\vec{a}_k}{\|\vec{a}_k\|}$.

### SI 2 Feasibility domain volume calculation

The volumetric modulus, or the normalized volume of a $P$-dimensional convex cone $\Omega$ is defined as the ratio

$$\Omega = \frac{\mathrm{vol}_P(\Omega \cap \mathbf{B}_P)}{\mathrm{vol}_P(\mathbf{B}_P)},$$

where $\mathbf{B}_P$) represents the closed unit ball in $\mathbf{R}^P$. This definition differs from the one given in [3] in that we consider the volume of the whole sphere rather than only a half space. Equivalently, the size of the feasibility domain can be quantified by the surface that it produces on a sphere. For example, for the unit sphere $\mathbf{S}_P$ in $\mathbf{R}^P$, it can be expressed as

$$\Omega = \frac{\text{vol}_{P-1}(\Omega \cap \mathbf{S}_P)}{\text{vol}_{P-1}(\mathbf{S}_P)},$$

and the intersection $\text{vol}_P(\Omega \cap \mathbf{S}_P)$ is called the cone's *solid angle* [3].

Due to the symmetry of the space, in general, the maximal size $\Omega$ of a convex cone is $\Omega = 1/2$, i.e., a half space. In our framework, because the parameters $\rho$ and $\mu$, arranged in the vector $b = [\rho \quad -\mu]$, must satisfy a sign condition, namely, $\rho > 0$, $\mu > 0$, the maximal size is further restricted to the positive orthant of the $P$-dimensional parameter space. Consequently, the maximal size of the feasibility domain here is $\Omega = 1/4$.

#### SI 2.1 Method to calculate the volume of the feasibility domain

For a minimal community of size $P = 2$, the volume of the feasibility domain can be explicitly calculated as

$$\Omega = \frac{1}{2\pi} \arccos(g_1 \cdot g_2),$$

which quantifies the proportion of the unit circle that is intersected by the generated cone. For a three-dimensional parameter space (the community is of size $P = 3$), the volume of the feasibility domain can also be explicitly calculated using the three linearly independent unit vectors $\{g_1, g_2, g_3\}$ as [6]

$$\Omega = \frac{1}{\pi} \arctan\left(\frac{|\det(g_1, g_2, g_3|)}{1 + g_1 \cdot g_2 + g_2 \cdot g_3 + g_1 \cdot g_3}\right).$$

The volume of a cone of arbitrary dimensions is given by [7]

$$\Omega = 2\frac{|\det G|}{\pi^{n/2}}\int_{\mathbb{R}^P_{>0}} e^{-\xi^\top G^\top G\xi} d\xi. \tag{3}$$

Setting $\frac{1}{2}\Sigma^{-1} = G^\top G$, the expression becomes the cumulative distribution of a multivariate normal distribution [5]

$$\Omega = \frac{2}{(2\pi)^{n/2}\sqrt{\det|\Sigma|}}\int_{\mathbb{R}^P_{>0}} e^{-\frac{1}{2}\xi^\top\Sigma^{-1}\xi} d\xi,$$

with the mean of zero and covariance matrix $\Sigma$. Note that (3) is equivalent to the expression that appears in [3]. This integral cannot be explicitly computed for most cases but can be approximated by numerical integration, multivariate series, or random methods [3]. Here we adopt a method to calculate the volume of the cone in which (3) is transformed into a cumulative distribution function of a multivariate normal distribution [8, 5]. In [9], several methods for the numerical computation of multivariate t-probabilities for hyper-rectangular integration regions are discussed, and Monte Carlo and quasi-Monte Carlo algorithms are provided. The authors argue that the latter improves the performance of the former. In our implementation, following [4, 5, 8], we use the R function `pmvnorm` which uses this method.

### SI 3 Simulations details

#### SI 3.1 Random communities generation, parameterization and simulations

We built random communities of sizes $S = (8, 6, 10, 12, 14, 16, 20)$, where $S$ is the total number of species and resources ($S = n + m$) for the CRM. Unless specified otherwise, we assume $n = m$, and the numerical values of the non-zero elements of the consumption matrix $C$ for the CRM, and the $A$ matrix for the competitive GLV were drawn using the absolute value of a normal distribution with mean $\eta_\text{o} = 0$ and standard deviation $\sigma = 0.6$, i.e., $|\mathcal{N}(\eta_\text{o}, \sigma)|$. The values of $\zeta_\beta$ were also drawn from the absolute value normal distribution $|\mathcal{N}(0, \sigma_\zeta)|$ with $\sigma_\zeta = 0.001$, for the case of resources' linear growth rate, and $\sigma_\zeta = \sigma$ for the case of logistic growth. To ensure the robustness of the results with respect to the parameterization, we also used uniform and Dirichlet distributions to draw random values for the numerical values of the system's parameters (SI 4 for more details). The connectance of the underlying network is calculated as $\kappa = \dfrac{C_\text{nz}}{nm}$, where $C_\text{nz}$ is the number of non-zero elements in $C$. For the simulations we generated 1000 communities of each size, with connectance $\kappa = \dfrac{n}{nm}$, and that satisfy $\text{rank}(C) = n$ i.e., every species consumes only one resource and every resource is only consumed by exactly one species, and calculated their feasibility volume. Then, we selected $\lfloor nm \rfloor$ zero elements from $C$, re-parameterized them drawing values from $\mathcal{N}(0, 0.6)$ and recalculated the feasibility domain. We iterated this procedure until $\kappa = 1$.

#### SI 3.2 NODF and rewiring

Using the same process for obtaining random communities described above, we built at least 500 of each size $n = m = (10, 15, 20)$, i.e., $S = (20, 30, 40)$, and connectance $\kappa = 0.2$ satisfying rank$(C) = n$. Then, we added nonzero elements to every community's $C$ matrix, such that it preserved its full rank and their connectance increased to $\kappa = (0.4, 0.6, 0.8)$. To modify the NODF of a generated community, we set a desired NODF value (NODF$_d$), and define a tolerance $tol = 0.05$ and constant $\bar{\lambda} = 0.5$, and run the following algorithm:

1. Calculate $\delta$ =abs(NODF($C$)-NODF$_d$). If $\delta > tol$ continue, else stop.

2. Randomly select a non-zero and a zero element of $C$ ($c_{\beta i} = \epsilon \neq 0$ and $c_{\alpha j} = 0$), and swap their numerical values such that $c_{\beta i} = 0$, $c_{\alpha j} = \epsilon$; call the resulting matrix $\bar{C}$.

3. If rank$(\bar{C}) < n$ go back to 1., else, make $\lambda = \bar{\lambda}$ and continue.

4. Calculate $\bar{\delta}$ =abs(NODF($\bar{C}$)-NODF$_d$). If $\delta - \bar{\delta} > 0$ continue, else, go to 2. with probability $1 - \lambda$ (continue with probability $\lambda$) and make $\bar{\lambda} = 0.5\lambda$.

5. Make $C = \bar{C}$ and go to 1.

#### SI 3.3 Niche overlap reparameterization

Assuming that the preferred interactions are on the main diagonal of $C$ for every community that we previously build (See SI 7 for details), we re-sampled this main diagonal drawing $n$ values from $\mathcal{N}(\eta_{\text{d}}, \sigma)$, where $\eta_{\text{d}} = 0$ and $\sigma = 0.6$ and singled out those communities that satisfied $S = 6$ and $\kappa = (0.55, 0.65, 0.8)$, $S = 10$ and $\kappa = (0.3, 0.9, 0.95)$, or $S = 20$ and $\kappa = (0.15, 0.5, 0.9)$. Because for this set of communities $\eta_{\text{d}} = \eta_{\text{o}} = 0$, they all have $\gamma = 1$. Then, we re-sampled their main diagonal four more times using

$\eta_{\rm d} = (1.53.04.56.0)$ obtaining $\gamma = (0.4, 0.25, 0.18, 0.14)$. After every main diagonal re-sampling, we calculated the feasibility volume of every community.

### SI 4 Robustness with respect to parameter distribution

To ensure that our results do not depend on the normal distribution of the numerical values of $C$, we used two additional distributions with biological significance (uniform and Dirichlet distributions) to generate random numerical values for the matrices' entries.

For the consumption matrix $C$, drawing random numbers from a normal distribution to assign numerical values to its non-zero entries (as in the results shown in the main text) represents the case when a few species have very small or very large consumption rates for a few resources, while most of the realized consumption interactions have a medium rate. When the non-zero elements of matrix $C$ are randomly drawn from a normal distribution, all consumption rates have the same probability of occurring. The parameterization process with uniform distributions is equivalent to the parameterization with a normal distribution (described in the Methods section): For every non-zero element in $C$, a single number is randomly drawn from $\mathcal{U}(m_C, M_C)$, respectively, where $0 > m_C > M_C$.

By using a Dirichlet distribution we simulate the case when all the species have the same overall consumption rate cap, and fractions of that rate are allocated to every resource that they consume [10]. To that end, to every non-zero element in column $i$ in $C$ we assign a numerical value from a vector generated by the function `rdiric` in the R package VGAM (with dimension equal to the number of the column $i$'s non-zero elements) such that $\sum_{\alpha=1}^{\alpha=m} c_{\alpha i} = Z_c$.

We repeated the simulations described in the Methods section building communities of sizes $P = (6, 10, 20)$ parameterizing the matrix $C$ using an uniform distribution with $m_C = 0.01$ and $M_C = 2\sigma$, and a Drichlet distribution with $Z_c = 2\sigma$. The feasibility volume $\Omega$ in both cases showed a decreasing trend as the connectance $\kappa$ increases. Also, using $\kappa P$ as a scaling variable for the volume of communities of various sizes resulted in a data collapse. Both results are qualitatively equivalent to those obtained using a normal distribution to parameterize $C$ (Figure S1).

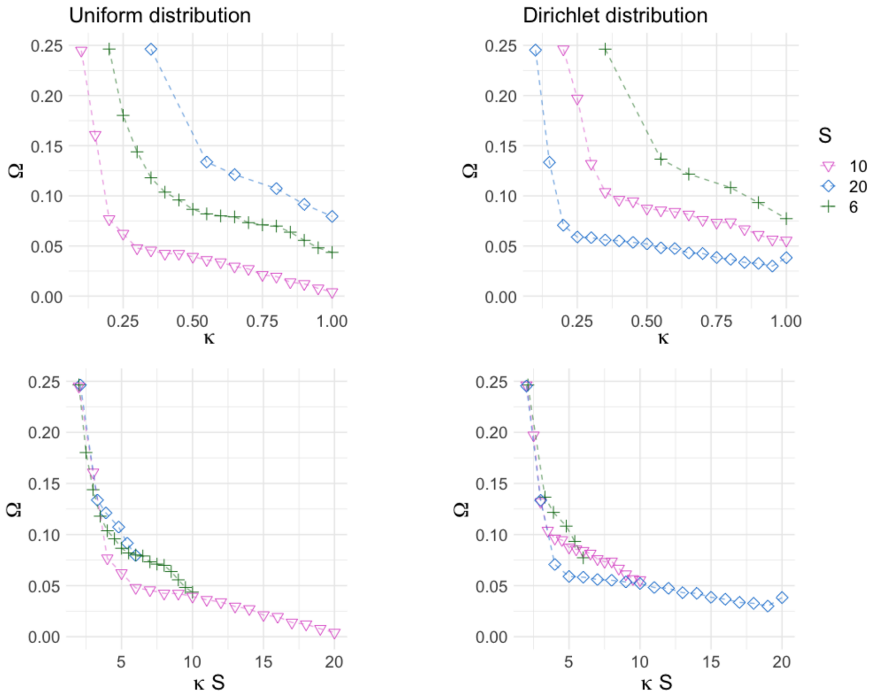

Figure S1: Parameterizing the consumption matrix $C$ using normal, uniform or Dirichlet distributions produced qualitatively equivalent results.

### SI 5 Scaling and power laws

Scaling laws are used to describe the relationship between two physical quantities that scale with each other. Mathematically, the phenomena can be explained by means of homogeneous functions which satisfy

$$f(sx) = s^k f(x),$$

for an integer $k$ and nonzero constant $s$, i.e., if the argument of a function is multiplied by a constant, its value is identical to the original one, multiplied by some power of that constant. Relationships of this kind have repeatedly appeared in expressions modeling different physical and biological phenomena [11]. It is commonly accepted that the existence of such relationships suggest that fundamental universal principles underlie much of the coarse-grained generic structure and organisation of living systems [12]. In biology and physics, body size and temperature are generally understood as determinants of variation when comparing different organisms or systems [12]. Thus, these two quantities are often used to investigate the existence of scaling laws in particular applications. We observed that the community size, when used as a scaling factor, leads to the data collapse in Figure 3 of the main text, which validates our choice.

A distribution of the form

$$y(x) = ax^b \tag{4}$$

for some $a > 0$ and $b > 0$, is said to follow a power law, and the plot of its log-log transformation, i.e., $\ln(y) = b\ln(x) + a$, is a straight line with slope $b$ and intercept $a$. The log-log plot of the distribution is also known as the complementary cumulative distribution function. This procedure constitutes a simple empirical test for whether a random variable has a power law.

We perform this test on the data obtained from our generated communities, i.e., we plot the log transformation of the product of the community's connectance $\kappa$ and its size $P$, v.s. the log transformation of the inverse of the feasibility volume. In other words, the former is the independent variable in (4), and the latter is the dependent one. The approximately straight lines in Figure S7 show that indeed, the relationship follows a power law. To estimate the numerical values of the constants $a$ and $b$, we fit the data using a linear regression which yields $a = 0.4956359$ and $b = 0.9942034$. The solid lines in Figure S8 show the resulting curves when the constants of the power law are approximated as $a = 1/2$ and $b = 1$, for the different community sizes, and the markers represent the data obtained from our generated communities.

### SI 6 Nestedness

Nestedness is a property of networks of species interactions that occurs when specialists tend to interact with proper subsets of the species that interact with more generalists [13]. Because nestedness is a concept that has only been verbally defined through the arrangement of interactions in communities, and not formally mathematically defined, there are several metrics, or methods that attempt to quantify it [14]. Here we use the nestedness metric based on overlap and decreasing fill (NODF), proposed by Almeida et al. [14], to quantify the nestedness of a community, which is based on the decreasing fill (or DF) and paired overlap (or PO) of the adjacency matrix. The DF compares the filling of adjacent rows (columns) in the community matrix through a quantity $\mathrm{DF}_{ij}$ ($\mathrm{DF}_{kl}$) for every pair of rows (columns) in the matrix where $j = i + 1$, ($l = k + 1$). The values, for the rows, are assigned as $\mathrm{DF}_{ij} = 100$ if $\mathrm{MT}_j \leq \mathrm{MT}_i$, where $\mathrm{MT}_i$ is the number of non-zero elements in row or column $i$, and $\mathrm{DF}_{ij}$=0 otherwise. The values of $\mathrm{DF}_{kl}$ (for the columns) are assigned analogously. The PO compares the zero-nonzero pattern of adjacent columns (rows) though quantities $\mathrm{PO}_{kl}$ ($\mathrm{PO}_{ij}$). For columns, $\mathrm{PO}_{kl}$ is the percentage of nonzero elements in column $l$ that are located at identical row positions to the nonzero elements in column $k$, and analogously for rows. For every pair of rows and columns, $\mathrm{N}_{pair}$ is defined as

$$\mathrm{N}_{pair} = \begin{cases} 0, & \mathrm{DF}_{pair} = 0 \\ \mathrm{PO}_{pair}, & \mathrm{DF}_{pair} = 100 \end{cases}$$

and the NODF is calculated as

$$\mathrm{NODF} = \frac{\sum \mathrm{N}_{pair}}{\frac{1}{2}\left(n(n-1) + m(m-1)\right)}.$$

Following this method, a perfectly nested community has a score NODF=1. For our simulations we converted the consumption matrix of our generated communities and measured their NODF using the function `nodf_cpp` in the *maxnodf* R package [15].

To vary the NODF in a community with a given connectance, we first obtained the maximum NODF and the corresponding structure of the $C$ matrix using the function `maxnodf` in the *maxnodf* R package. Then, we parameterized the non-zero elements of $C$ drawing numbers from $\mathcal{N}(o, \sigma)$ and taking the absolute values, and we implemented the following algorithm that rewires the community network to decrease the NODF, preserving its connectance, ensuring that the resulting matrix fulfills the full rank condition.

1. Calculate NODFini = NODF($C$).

2. Randomly select a non-zero and a zero element of $C$ ($c_{\beta i} = \epsilon \neq 0$ and $c_{\alpha j} = 0$), and swap their numerical values such that $c_{\beta i} = 0$, $c_{\alpha j} = \epsilon$; call the resulting matrix $\bar{C}$.

3. If $\text{rank}(\bar{C}) < n$ go back to 2., else, calculate NODFs = NODF($\bar{C}$) and continue.

4. If NODFini - NODFs > 0, make $C = \bar{C}$ and go to 1.

#### SI 6.1 NODF normalization and control for modifiers

NODF is known to be highly correlated with the size and connectance of the community which, in addition to the maximum and minimum possible values for networks with some specific connectances being far from the [0,1] boundaries (Fig S2), could make the comparison of nestedness across networks solely using NODF troublesome [15]. To overcome this issue, Song et al. [15] propose the normalization $\text{NODF}_\text{n} = \dfrac{\text{NODF}}{\max(\text{NODF})}$, which can then be corrected for connectance and community size as $\text{NODF}_\text{c} = \dfrac{\text{NODF}_\text{n}}{\kappa \log\left(\overline{[n, m]}\right)}$ , where $\overline{[\cdot]}$ denotes the geometric mean. We use this normalization to compare the nestedness measurement across our communities of different sizes and connectances.

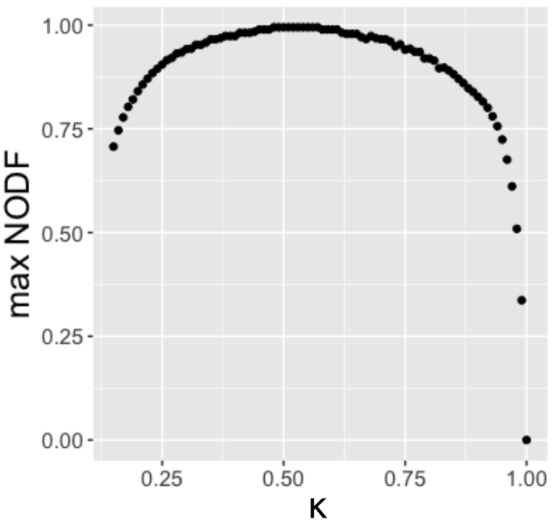

Figure S2: The NODF can take values between 0 and 1. However, for networks with small or large connectance, the realizable maximum is below 1.

### SI 7 Niche overlap

To vary the niche overlap of a community, we re-sampled the consumption rates between every species and its preferred resource. Because the $C$ matrix has to be of full rank, for a community with $n = m$, $n$ unique pairs of interacting species and resources can be found. Each of these unique pairs contains a single species and a resource that it consumes (their preferred resource), and the resource doesn't appear in any of the other pairs (left panel in Figure S3). The The $C$ matrix can then be reordered such that the interactions between the unique pairs are on its main diagonal (center and right panels in Figure S3). Thus, we consider that the diagonal interactions represent every species' consumption rate of its exclusive preferred resource.

To re-sample the pairs' consumption rates, we drew numbers from a normal distribution with the mean of $\eta_{\mathrm{d}} \geq \eta_{\mathrm{o}}$ and standard deviation $\sigma$, and then taking the absolute value. Recall that all the non-zero interaction strengths of $C$ (now the off-diagonal elements) have been previously parameterized using the absolute value of a normal distribution with the mean of $\eta_{\mathrm{o}}$ and standard deviation $\sigma$. The parametric similarity between the diagonal and the off-diagonal interactions in the $C$ matrix, is quantified for a general consumption matrix by

$$\gamma = \frac{\eta_{\mathrm{o}} + 1}{\eta_{\mathrm{d}} + 1}.$$

Because we used $\eta_{\mathrm{o}} = 0$ to parameterize the $C$ matrix in the main text, the above expression simplifies to

$$\gamma = \frac{1}{\eta_{\mathrm{d}} + 1}.$$

Figure S3.b shows the decrease in feasibility volume in communities of size $S = (6, 10, 20)$

as their connectance increases, for values of $\gamma = (1, 0.17, 0.09)$. Note that all the communities used for the other results have $\gamma = 1$ because they were parameterized by drawing all the non-zero interaction strengths from only one normal distribution, i.e., $\eta_\text{o} = \eta_\text{d}$ , and the resources can always be reordered such that the main diagonal has only non-zero elements.

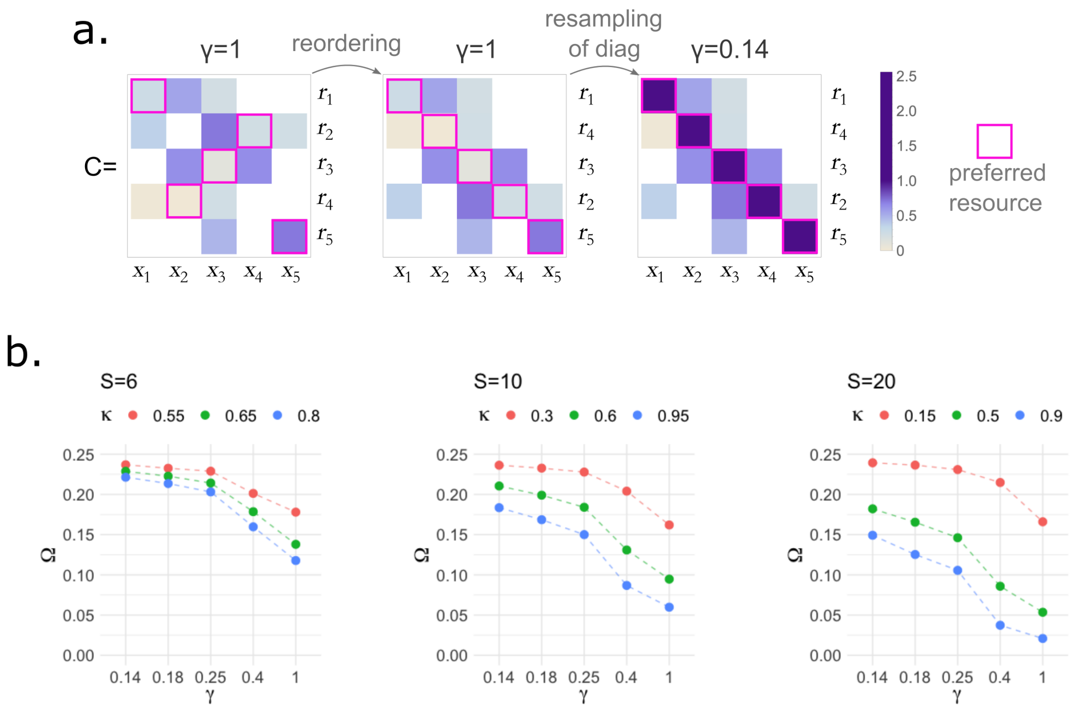

Figure S3: A greater niche overlap leads to a loss of feasibility. A greater niche overlap occurs when competition between species for specific resources increases. **a.** We modify the niche overlap in our synthetic communities by re-sampling the rate at which every species consumes its exclusive preferred resource. For a community of size $S = 10$ ($n = m = 5$), the left panel shows that the preferred exclusive resource for species $x_1$, $x_2$, $x_3$, $x_4$ and $x_5$ are $r_1$, $r_4$, $r_3$, $r_2$, and $r_5$, respectively. The center panel shows a reordering of the resources such that the interactions with the species' preferred resource is on the main diagonal. Here, the off-diagonal and diagonal elements of $C$ have been randomly sampled using the absolute value of a normal distribution with the mean of $\eta_{\rm o}$ and $\eta_{\rm d} = \eta_{\rm o}$, respectively, and standard deviation $\sigma$. The right panel shows the consumption matrix when the interactions between the species and their preferred resource have been re-sampled using the absolute value of a normal distribution with mean $\eta_{\rm d} \geq \eta_{\rm o}$ and standard distribution $\sigma$. We quantify the niche overlap as $\gamma = (\eta_{\rm o} + 1)/(\eta_{\rm d} + 1)$. Because we consider values $\eta_{\rm d} \geq \eta_{\rm o}$, $\gamma$ can take values between 1 and 0; $\gamma \approx 0$, indicates that all species are specialists that consume exclusive resources (no niche overlap), and $\gamma = 1$ indicates that the consumption preferences are evenly distributed across the consumption matrix (and any pair of species can compete for any resource). **b.** The panels show the feasibility volume of random communities with respect to their $\gamma$ value (horizontal axis), for different connectances (colored markers), in communities of size $S = (6, 10, 20)$. We find that for a given connectance, a stronger niche overlap tends to shrink the feasibility volume across the different community sizes. This suggests that parametric dissimilarity, which reduces the niche overlap between species, thus contributes to a higher likelihood of coexistence.

### SI 8 Additional Figures

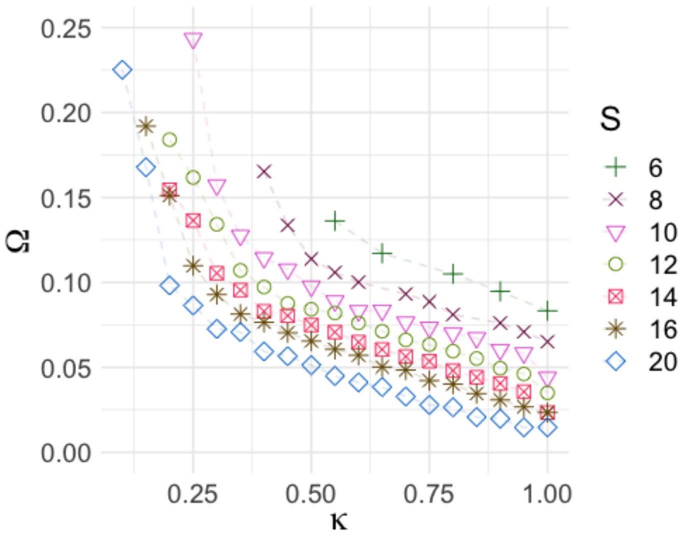

Figure S4: Volume of the feasibility domain of communities of various sizes with respect to their connectance $\kappa$. The markers represent the mean volume of 1000 communities. Dotted lines are added to enhance readability.

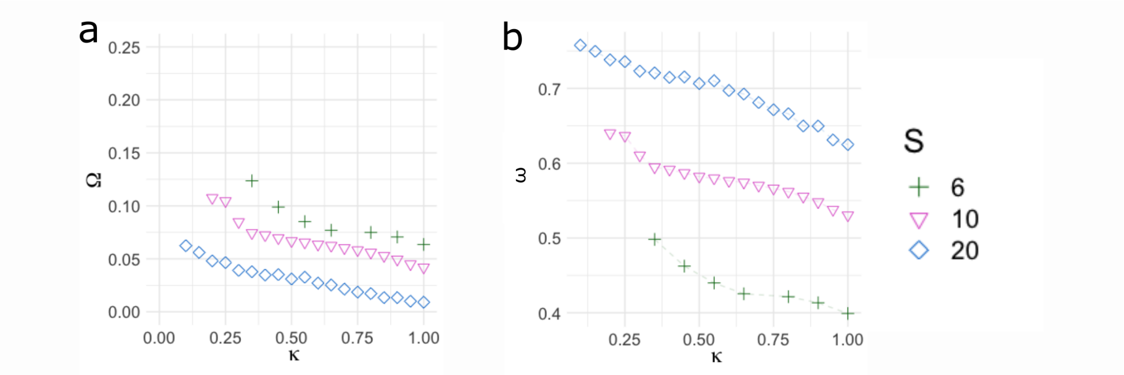

Figure S5: When the resources grow at a logistic rate, i.e., the non-zero values of $\zeta$ are in the same order of magnitude as those of $C$, the feasibility connectance trends are not qualitatively modified, but the feasibility of the whole community decreases at low connectances of $C$. For the simulations we parameterized $\zeta_{\beta\beta}$ by drawing random values from a normal distribution $\mathcal{N}(0, \sigma)$ and then taking the absolute value. **a.** Feasibility of the whole community with respect to the system's connectance and size. **b.** Feasibility per species with respect to the system's connectance and size

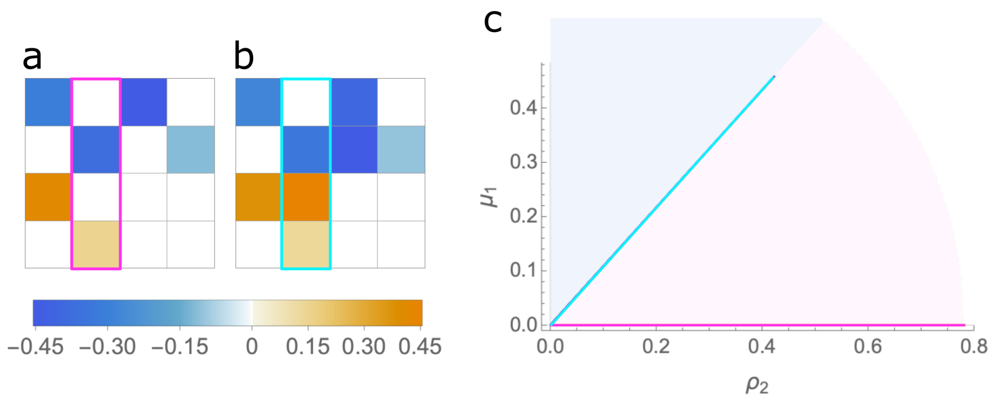

Figure S6: A more connected consumption matrix tends to lead to a shrinkage of the feasibility domain. Consider a consumer resource community with 2 species and 2 resources. **a**. The panel shows one example of $A$ matrix resulting from writing the system in the GLV form. The matrix can be divided into four 2×2 blocks: the upper left represents $-\zeta$, the upper right represents $-C$, and the lower left represents $\theta C^\top$ . Here $C$ is a diagonal matrix, i.e. every species consumes one exclusive resource. **b**. Panel shows a different $A$ matrix for the community with size $S = 4$. Here, $C$ contains one extra interaction with respect to panel **a**, and all the rest of the parameters are identical. **c**. 2D projection on the $(\rho_2, \mu_1)$ plane of one generating vector from **a** and **b**, respectively. The pink vector, coming from the matrix with diagonal $C$, lies in the $\rho_2$ axis, indicating that any value of $\rho_2$ and $\mu_1$ will lead to a feasible equilibrium. The cyan vector, coming from the matrix with an added consumption interaction cuts the $(\rho_2, \mu_1)$ plane, confining the feasible parameter combinations to the blue area.

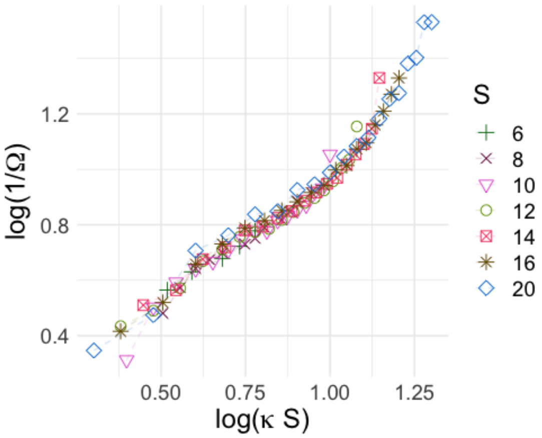

Figure S7: Log transformation of the product of the community's connectance $\kappa$ and its size $P$, v.s. the log transformation of the inverse of the feasibility volume. The approximately straight line shows that the relationship between the two variables follows a power law.

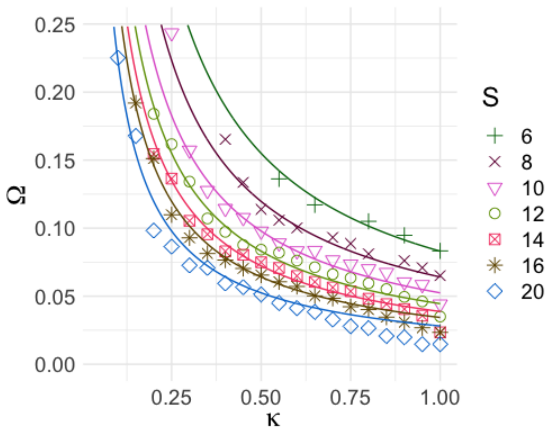

Figure S8: The relationship that describes the decrease in volume of the feasibility domain when the complexity of the system increase is $\Omega = \dfrac{1}{2\kappa S}$. Solid lines in the panels show the resulting curve over the computed volumes of communities of various sizes (shown by the markers).

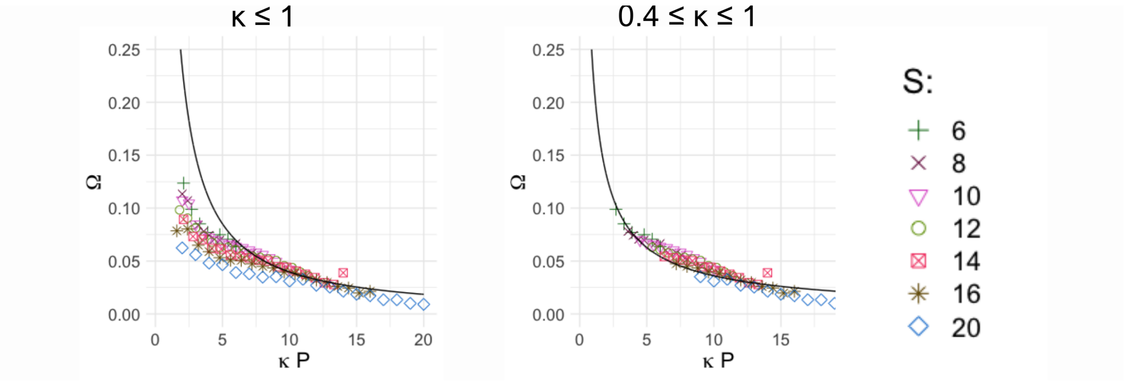

Figure S9: The power law to describe the complexity-feasibility relationship does not trace the simulation values when for the case of resources' logistic growth, particularly at low values of $\kappa S$ (left panel). However, if only more connected communities are considered ($\kappa \geq 0.4$), the complexity-feasibility relationship, modeled by a power law, is recovered.

### SI 9 References
